# Large-scale structure prediction by improved contact predictions and model quality assessment

**DOI:** 10.1101/128231

**Authors:** Mirco Michel, David Menéndez Hurtado, Karolis Uziela, Arne Elofsson

**Affiliations:** Science for Life Laboratory and Department of Biochemistry and Biophysics, Stockholm University, Stockholm 10691, Sweden

## Abstract

**Motivation:** Accurate contact predictions can be used for predicting the structure of proteins. Until recently these methods were limited to very big protein families, decreasing their utility. However, recent progress by combining direct coupling analysis with machine learning methods has made it possible to predict accurate contact maps for smaller families. To what extent these predictions can be used to produce accurate models of the families is not known.

**Results:** We present the PconsFold2 pipeline that uses contact predictions from PconsC3, the CONFOLD folding algorithm and model quality estimations to predict the structure of a protein. We show that the model quality estimation significantly increases the number of models that reliably can be identified. Finally, we apply PconsFold2 to 6379 Pfam families of unknown structure and find that PconsFold2 can, with an estimated 90% specificity, predict the structure of up to 558 Pfam families of unknown structure. Out of these 415 have not been reported before.

**Availability:** Datasets as well as models of all the 558 Pfam families are available at http://c3.pcons.net/. All programs used here are freely available.

**Supplementary information:** No supplementary data

## 1 Introduction

A few years ago maximum entropy methods revolutionized the accuracy of contact predictions in proteins (Weigt *et al*., 2009; Burger and van Nimwegen, 2010; Aurell, 2016). This enabled the prediction of accurate protein models using no information from homologous protein structures (Marks *et al*., 2011; Morcos *et al*., 2011). It has been shown that accurate protein structures can be obtained for soluble proteins (Marks *et al*., 2011), membrane proteins (Nugent and Jones, 2012; Hopf *et al*., 2012; Hayat *et al*., 2015) and even disordered proteins (Toth-Petroczy *et al*., 2016). These methods have also been used to predict interactions between proteins (Weigt *et al*., 2009; Ovchinnikov *et al*., 2014; Hopf *et al*., 2014).

Until recently such methods have been limited to very large protein families (Kamisetty *et al*., 2013; Skwark *et al*., 2014). However, by the inclusion of additional information and improved machine learning methods it is now often possible to obtain accurate contact maps for families as small as a few hundred effective sequences (Michel *et al*., 2017; Jones *et al*., 2015; Wang *et al*., 2017).

Pfam contains today approximately 16,000 protein families that vary in size between a few tens to hundreds of thousands effective sequences. About half (46%) of these protein families contain no representative structure, i.e. there is more than 7,500 protein families without a structure. The families with structure are on average larger than the ones without, median size 680 vs. 134 effective sequences, i.e. most of the families without a structure are too small for maximum entropy contact prediction but might be within reach for methods that combine DCA and advanced machine learning.

Now, we ask the question how many of these roughly 7,500 protein families without a structure can be modeled reliably by using state of the art contact prediction methods. To the best of our knowledge the largest effort to model protein families was performed by the Baker group who modeled structures for 614 families by including a very large set of sequences from meta-genomics (Ovchinnikov *et al*., 2017). However, their approach for contact prediction was based on a maximum entropy method (Antala *et al*., 2015) and not the newer methods using machine learning.

The PconsFold2 pipeline is described in Figure 1. Given an input sequence PconsFold2 generates four alignments. These alignments are then used by PconsC3 (Michel *et al*., 2017) to predict four different contact maps. The 2.5L (L=length of sequence) top ranked contacts are then used to fold the protein. In contrast to PconsFold (Michel *et al*., 2014) PconsFold2 uses CONFOLD (Adhikari *et al*., 2015), i.e. the NMR protocol of CNS (Brunger, 2007) and not ROSETTA (Leaver-Fay *et al*., 2011) to generate 50 models for each contact map. This makes the pipeline much faster but possibly slightly less accurate.

**Fig. 1.**
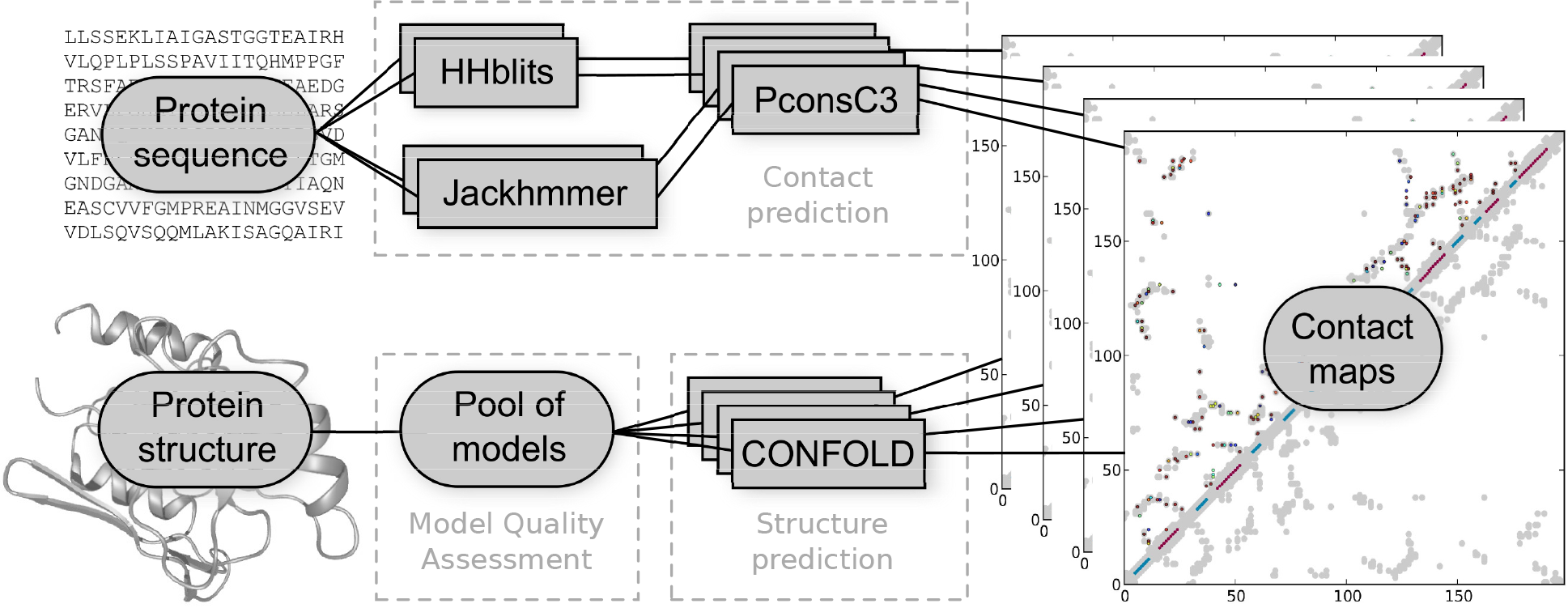
PconsFold2 workflow. Given an input sequence four alignments are created using HHblits and Jackhmmer at two different E-value thresholds of 1 and 10^−4^. Based on these alignments PconsC3 generates four contact maps. The 2.5L (L=length of sequence) top ranked contacts are then used by CONFOLD to generate 50 models for each alignment, resulting in 200 models for each query sequence. These models are finally ranked by a model quality assessment program.

The final step in the pipeline is the model quality assessment. In addition to existing model quality assessment methods we introduce PcombC. PcombC is a linear combination of three separate assessment scores: Pcons (Lundström *et al*., 2001), ProQ3D (Uziela *et al*., 2017), and the agreement between predicted and observed contacts in the model.

Finally, we apply PconsFold2 to 6379 Pfam families without a known structure. Using the cutoffs for a false positive rate of 10% we do identify 558 protein families where the first ranked model is estimated to be correct by one of the three model quality estimators. Out of these 558 Pfam families 415 are not present in the earlier studies by the Baker group (Ovchinnikov *et al*., 2017, 2015).

## 2 Methods

### 2.1 Datasets

There are 16,295 protein domain families in Pfam 29.0. Out of these 7733 domains have a known structure with a HHsearch (Söding, 2005) hit in PDB with an E-value of less than 10^−3^ that covers at least 75% of it’s representative sequence. For each Pfam family we assign a representative sequence. We refer to the length and properties of a Pfam family by the representative sequence. The representative sequence of a Pfam domain with known structure is set to be the protein sequence ranked first by HHsearch against the PDB database bundled with HHsuite (Meier and Söding, 2015) (date: 2016-09-07)

The test dataset was generated from 626 Pfam domains that were randomly selected from 6925 domains with known structure that are longer than 50 residues.

From the remaining Pfam domains we excluded all Pfam domains that can be found in the `pdbmap` file from Pfam release 29.0 and those shorter than 50 residues. This results in a set of 7537 Pfam domains with unknown structure. For each of these sequence we define the highest ranked sequence in the HHblits (Remmert *et al*., 2012) alignment against uniref20 (date: 2016-02-26) to be the reference sequence of the family.

### 2.2 Alignments

The input to direct coupling analysis (DCA)-based contact prediction methods is a multiple sequence alignment. These alignments were generated using both HHblits and Jackhmmer (Eddy, 2011), each at Evalue thresholds of 1 and 10^−4^. HHblits was run against the uniprot20 database from HHsuite (date: 2016-02-26). The parameter `-all` has been used and `-maxfilt` and `-realign_max` were set to 999999 as in (Michel *et al*., 2017). Jackhmmer searches were performed against Uniprot90 (Magrane and Consortium, 2011) (2016-04-13) and were run for five iterations with both -E and -incE set to the respective E-value cutoffs. The searches were started from the Pfam representative sequence, i.e. the Pfam alignments were ignored. Alignment (family) size is measured in effective sequences (M_eff_) as defined in (Ekeberg *et al*., 2013).

### 2.3 Contact prediction

PconsC3 is used to predict contacts between pairs of amino-acids in the Pfam reference sequences. To overcome the limit of DCA methods requiring large alignments, PconsC3 combines the results of such methods with contacts predicted by a machine-learning based method (Michel *et al*., 2017). It then uses a similar pattern recognition approach as PconsC2 (Skwark *et al*., 2014) to iteratively increase the quality of the predicted contact map. PconsC3 was run as described earlier (Michel *et al*., 2017). However, PconsFold2 uses all four alignments as inputs predicting one contact map for each alignment. Contact map quality is measured in positive predictive value (PPV) over the same number of top-ranked contacts that were used during folding (2.5 · sequence length (L)). It should be noted that most earlier papers report the PPV for L or even fewer contacts. The average contact score for a contact map refers to the mean PconsC3 score of these 2.5 L contacts.

### 2.4 Model generation

Contacts predicted by PconsC3 are then applied as distance restraints between the C*ß*-atoms (C*α* in the case of glycine) during protein structure prediction. We use CONFOLD (Adhikari *et al*., 2015) for this task. Secondary structure predictions from PSIPRED (Jones, 1999) are also used as inputs. When folding a protein using CONFOLD a fixed number of contacts are used. Here, contacts are sorted by their PconsC3 score and a threshold is set on the number of top-ranked contacts to use. This threshold is based on the length of the sequence as an input to CONFOLD, which folds the protein using CNS (Brunger, 2007). It is to be noted that we only run the first stage of CONFOLD and omit the refining second stage in order to keep runtime low. For each alignment we generate 50 models resulting in a pool of 200 models per Pfam family.

### 2.5 Model ranking

The CONFOLD pipeline generates several models that by default are ranked by the CNS contact energy (NOE). This is the sum of all violations of all contact restraints used to generate a model. In order to make this score comparable between targets we normalize it by the length of the input protein sequence and refer to it as CNS-contact.

In addition to using the CNS contact energy we also used ProQ3D (Uziela *et al*., 2017) and Pcons (Lundström *et al*., 2001). Further we developed PcombC a linear combination between the scores of ProQ3D, Pcons, and PPV, similar to what we used in CASP4 (Wallner *et al*., 2003) and CASP5 (Wallner and Elofsson, 2005a). Coefficients have been determined using a grid-search on a 10x10x10 grid with values ranging from 0 to 1 and a step size of 0.1, optimizing the area under the ROC-curve for determining whether a model is correct or not (TM-score threshold of 0.5). In order for the score to remain within the same scale as the input scores, the coefficients have been normalized to:

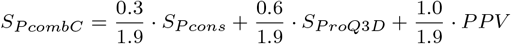

### 2.6 Evaluation

Model quality is measured in template modeling score (TM-score) scores (Zhang and Skolnick, 2004). For the ROC-analysis we set a TM-score threshold of 0.5 to distinguish between correct and incorrect models.

### 2.7 Runtime

The running time of the folding step was measured on a single core of an Intel Xeon E5-2690 v4 processor. For the test dataset it takes around 30 seconds on average to generate one model with a minimum of 4s per model for the shortest family (50 residues) and 245s for the longest (524 residues).

## 3 Results

### 3.1 Utilization of predicted contacts

First we set out to find the best way to generate models using contacts predicted from PconsC3 (Michel *et al*., 2017) and the CONFOLD (Adhikari *et al*., 2015) folding algorithm. Preliminary data indicated that using a threshold for the number of contacts utilized during folding of 2.5 times the length of the sequence is close to optimal. Using this threshold we then investigated the effect of the number of generated models on the quality of the best and top-ranked model. Here, the CNS-contact score from CONFOLD is used to rank the models. Increasing the number of generated models from the default 20 to 50 does not increase the quality of top-ranked models (average TM-score 0.40). However, the average TM-score of the best among all generated models increases from 0.43 to 0.45. Increasing the number of models further (up to 200) only generates a marginal improvement of the best TM-score to 0.46. We thus decided that generating 50 models for a given contact map is a good tradeoff between model qualities and running time.

It has previously been observed that the quality of predicted contacts depends on the underlying alignment (Skwark *et al*., 2013). We therefore tried to identify the optimal alignment method and E-value cutoff. In addition we investigated whether model quality can be improved by using a set of alignments with varying methods and E-value thresholds instead of a single fixed alignment. In figure 2 it can be seen that the performance is similar for all four alignment methods used both regarding the agreement with the contact map and the average TM-score of the generated models, see Table 1. However, the quality of both the top-ranked (0.42) and best possible model (0.49) is improved slightly when using a combination of all alignments. Therefore we decided to use this for the pipeline.

**Fig. 2.**
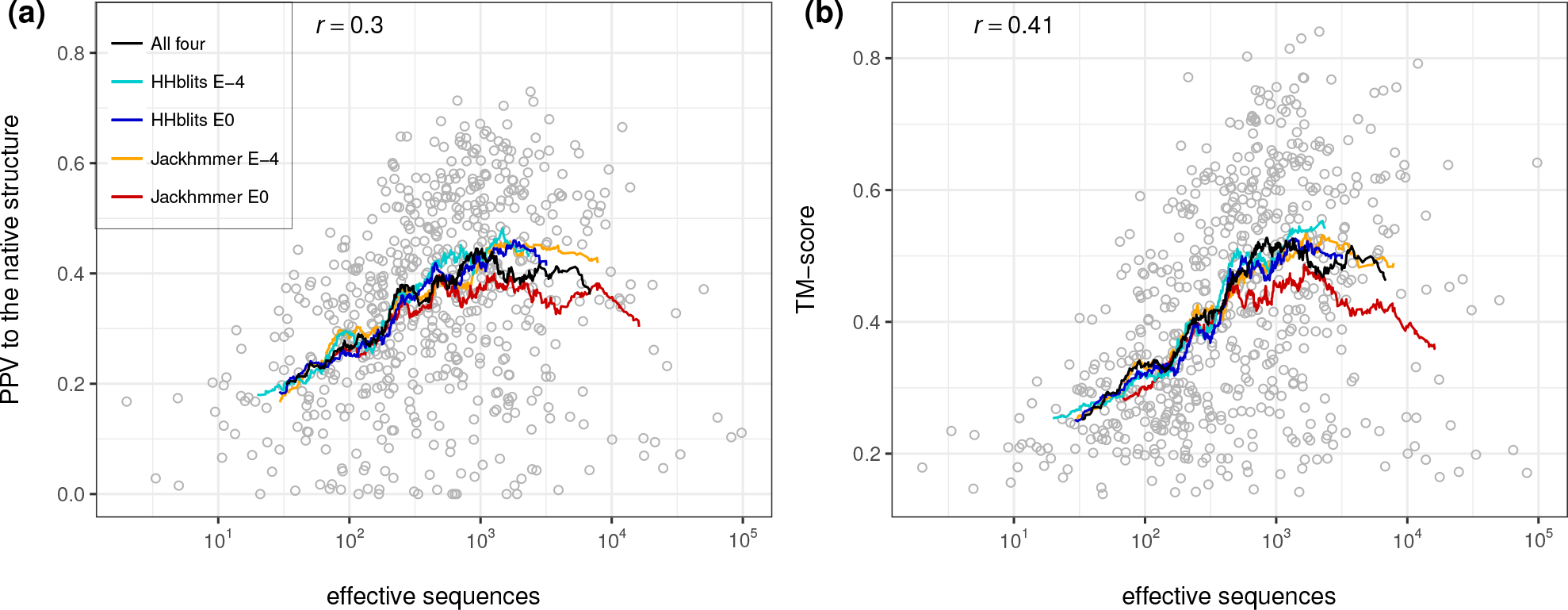
Family size in effective sequences of the benchmark dataset against (a) PconsFold2 model quality in TM-score of the top-ranked models and (b) against contact map quality in PPV. Lines show moving averages for using the HHblits alignment at E-value 10^−^ 4 (cyan), HHblits at E-value 1 (blue), Jackhmmer at E-value 10^−^ 4 (orange), Jackhmmer at E-value 1 (red), and all four alignments combined (black). Circles show individual results when using all four alignments combined.

**Table 1.** Performance for different alignment methods.

|  | PPV to native | TM-score (top) | TM-score (max) |
| --- | --- | --- | --- |
| All | 0.35 | 0.42 | 0.49 |
| HHblits $10^{-4}$ | 0.34 | 0.40 | 0.45 |
| HHblits $10^0$ | 0.34 | 0.40 | 0.45 |
| Jackhmmer $10^{-4}$ | 0.35 | 0.41 | 0.46 |
| Jackhmmer $10^0$ | 0.33 | 0.40 | 0.44 |

### 3.2 Overall performance

Figure 2 shows the performance of the pipeline of the top-ranked model against the family size, measured in effective sequences of the underlying alignment. Here, models were ranked by their CNS-contact score as described above. Generally, both contact prediction accuracy and model accuracy clearly depends on family size with correlations of 0.30 for PPV and of 0.41 for TM-score. However, what might be most notable is that there is a large variation between different proteins. Some protein families with as few as 100 effective sequences both have good contact predictions and good final models, while some proteins with 10^5^ are no better than completely random models. This indicates that we need to use other evaluation procedures to identify the correct models.

Table 2 further reveals that for families smaller than 100 effective sequences, structure prediction has a success rate (percentage of models that are above 0.5 TM-score) of only 3%. As soon as the family becomes larger than 100 effective sequences the number of correct models increases rapidly. The success rate is 34% for families between 100 and 1000 effective sequences and 52% for large families with more than 1000 effective members.

**Table 2.** Fraction of correct models and contact maps.

| $M_{\text{eff}}$ | >0.5 PPV | >0.5 TM-score |
| --- | --- | --- |
| <100 | 0% | 3% |
| 100-1000 | 22% | 34% |
| >1000 | 35% | 51% |

### 3.3 All contact maps do not generate good models

Next we compared the quality of the models (TM-score) with the accuracy of the predicted contacts (PPV), Figure 3a. Clearly, there is a strong Pearson correlation (*r*=0.63). There is also a weaker correlation (*r*=0.45) between the strength of the PconsC3 contact prediction and the quality of the model, Figure 3f.

**Fig. 3.**
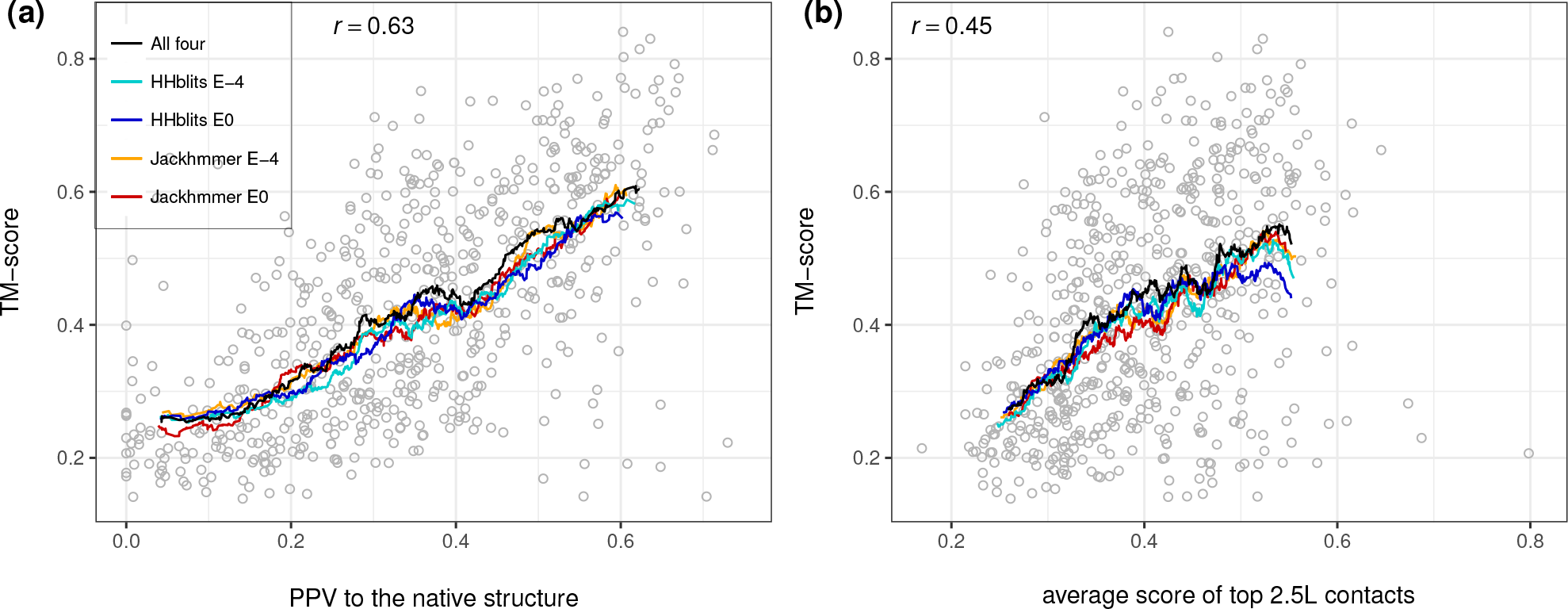
TM-score of the models using CONFOLD and CNS-contact ranking against (a) PPV of the underlying 2.5L contacts to the native structure and (b) the average score of these contacts.

From Figure 3 it is clear that there exist a number of bad models for large families with good contacts and also good models for small families with weakly predicted contacts. This indicates that it is not sufficient to only use the contact map or number of sequences in the alignment for identification of the successfully predicted protein models. Instead it might be better to evaluate the models directly.

### 3.4 Model quality assessment

To estimate the quality of a protein model model quality assessment methods can be used. Model quality estimators can be divided in consensus and single model methods. Here, we have used the Pcons consensus method to predict a quality score for each model based on a comparison of all models against each other (consensus method) (Wallner and Elofsson, 2005b). For single model quality estimation we have used ProQ3D a method that assesses the quality of a single model using deep learning (Uziela *et al*., 2017). Both of these methods have been shown to perform among the best methods in CASP.

It is also possible to evaluate the quality of a model by comparing how well it satisfies the predicted contacts. This can be done by either just counting the number of fulfilled contacts (PPV) or using the CNS-NOE energy. Both these methods perform very similar in terms of this evaluation (data not shown). We thus only report CNS-contact in all further analysis.

We have also developed a combined model quality estimator, PcombC, using all three methods. PcombC is a linear combination between the scores of these three methods, similar to what we used in CASP4 (Wallner *et al*., 2003) and CASP5 (Wallner and Elofsson, 2005a). PcombC has been optimized for discriminating between accurate and inaccurate models. This has the advantage of being able to interpret the predicted score in terms of absolute model quality, enabling statements about the confidence of a predicted model being correct.

Figure 4 shows how the scores of different QA tools predict TM-score. As before, for each family in the test set we ranked all models by each QA score and selected the top ranked model. Pearson correlation (r) is highest for PcombC, followed by Pcons and ProQ3D, while the CNS-contact energy correlates slightly worse. However, the average TM-score of the top-ranked model is almost identical for all four methods.

**Fig. 4.**
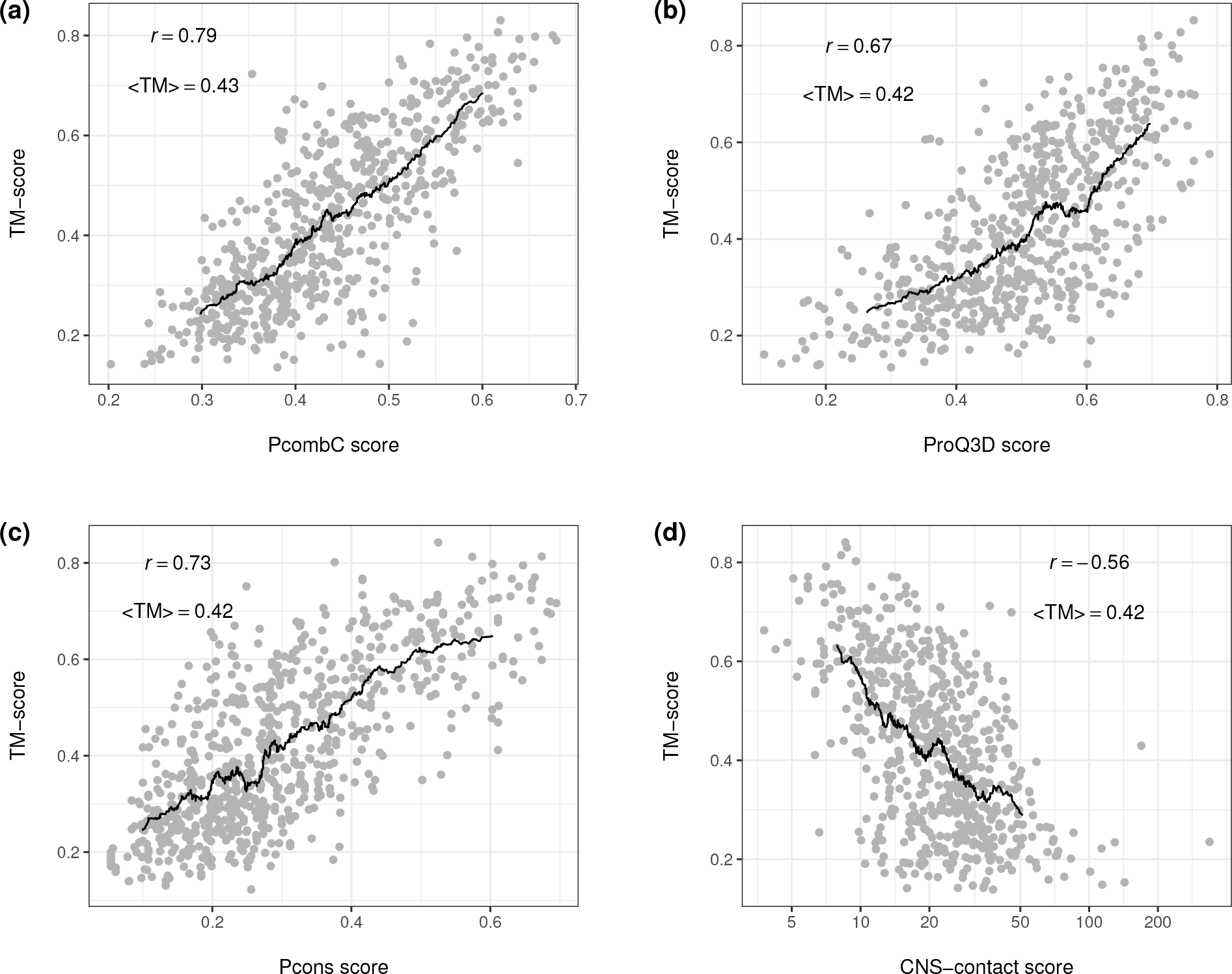
Model quality in TM-score on the benchmark dataset of the top-ranked models against (a) PcombC score (scoring function of the PconsFold2 pipeline), (b) ProQ3D score, (c) Pcons score, (d) and (d) CNS contact energy normalized by sequence length. Pearson correlations *r* and average TM-scores (<TM>) are shown, black lines represent moving averages with a window of 60 proteins. For CNS-contact the Pearson correlation has been calculated on log10(CNS-contact)

### 3.5 ROC-curve

In order to estimate model quality when there is no known structure available it needs to be predicted as accurately as possible. The goal is not only to select the best models from a set of predictions but also to predict how much these models can be trusted. Figure 5 shows the false positive rate (FPR) for different quality assessment (QA) tools when classifying predictions into correct (TM-score ≥ 0.5) or incorrect (TM-score < 0.5) models.

**Fig. 5.**
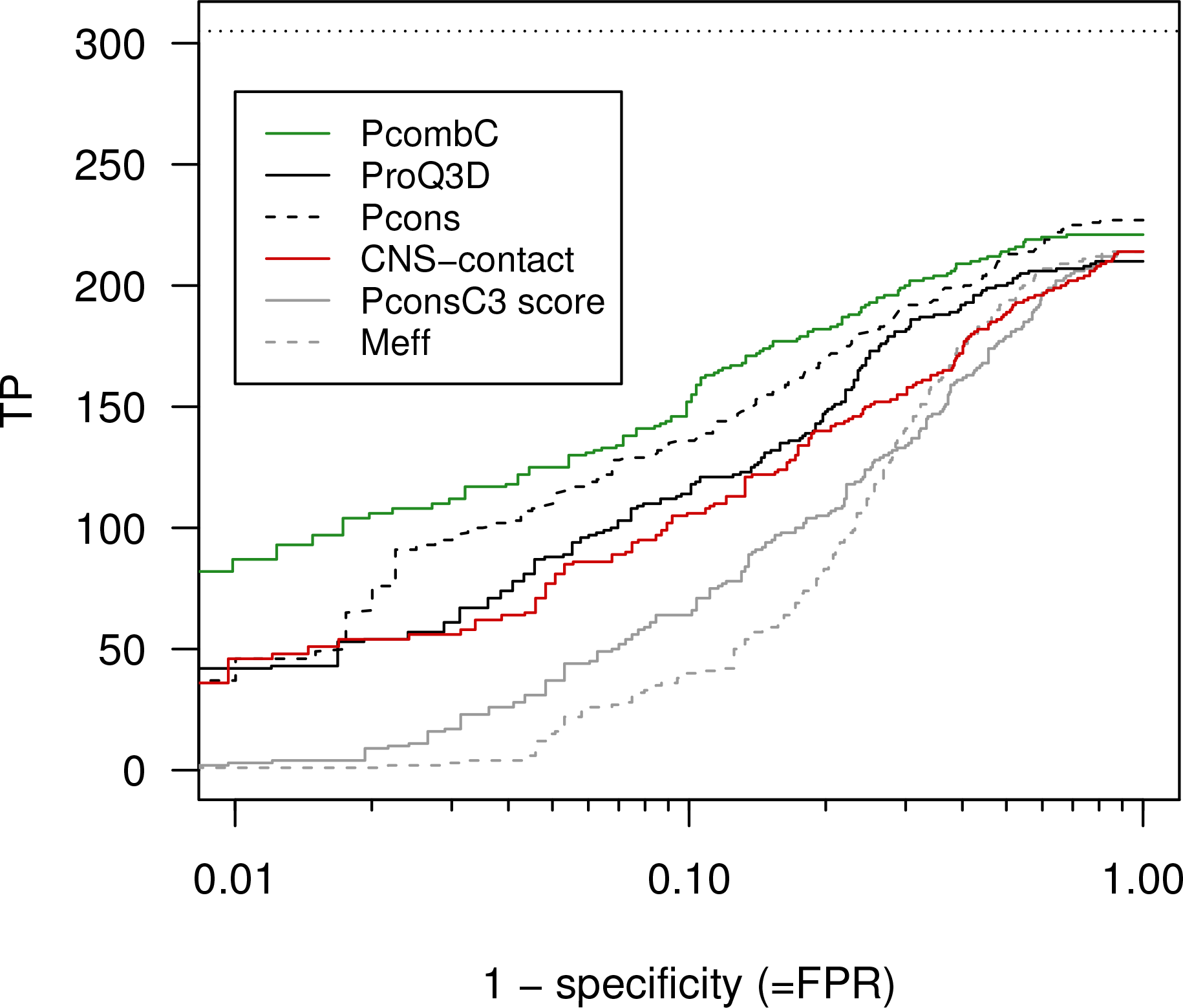
ROC-like plot for different ways of evaluating and ranking the models in the benchmark dataset. While the x-axis shows true positive rate on a logarithmic scale, the y-axis shows the number of proteins with TM-score ≥ 0.5. The horizontal line indicates the best possible outcome, i.e. the number of families with TM-score≥ 0.5 when ranking the models by TM-score score.

Although the overall number of correct top-ranked models does not change much (number of true positives at FPR = 1.0), there are differences in the ability of the different scores to classify the models. It can be seen that all methods evaluating the model quality are significantly better than just using the number of effective sequences or the PconsC3 scores, see Figure 5.

Table 3 also shows that the combined method, PcombC, clearly is better at identifying methods than any of the individual methods. At a FPR of 10% PcombC identifies 152 models compared with 106-136 for the individual methods. At the more strict 1% FPR PcombC identifies 29% of the models compared with 12-15% for CNS-contact and ProQ3D and Pcons.

**Table 3.** ROC analysis when classifying whether a model is correct (TM-score ≥ 0.5) or not.

| Method | # models at FPR |  |  |
| --- | --- | --- | --- |
|  | 0.01 | 0.1 | 1.0 |
| PcombC | 87 (29%) | 152 (50%) | 221 (72%) |
| Pcons | 37 (12%) | 136 (45%) | 227 (74%) |
| ProQ3D | 42 (14%) | 114 (37%) | 210 (69%) |
| CNS-contact | 46 (15%) | 106 (35%) | 214 (70%) |
| PconsC3 score | 3 (1%) | 64 (21%) | 214 (70%) |
| $M_{\text{eff}}$ | 1 (0.3%) | 40 (13%) | 214 (70%) |
| best possible | 305 | 305 | 305 |

The results shown here indicate that a combination of QA methods along with contact map agreement provides a significant improvement in detecting correct models over the best single methods. It can be used to reliably predict model accuracy while at the same time being more sensitive than previous methods.

## 4 Discussion

We then applied PconsFold2 as well as all quality assessment tools to the set of Pfam domains without known structure. This resulted in predicted structures for 6379 Pfam families (85% of all Pfam domains with unknown structure) 4. Based on the results from the ROC analysis on the test dataset we set the score cutoffs at FPR 0.01 and 0.1. We then use these cutoffs to estimate how many Pfam families of unknown structure can be predicted accurately (TM-score ≥0.5) at a given FPR. The union is defined as the non-overlapping number of families for which any quality assessment method predicts a model to be accurate.

At 0.01 FPR, or 99% specificity, models for a total of 114 Pfam families are predicted to be accurate, Table 4. This number increases almost four fold to 558 families, when allowing for a FPR of 0.1. More than 74% of these families do not overlap with other large scale structure prediction studies (Ovchinnikov *et al*., 2017, 2015). This indicates that our approach of using improved contact prediction combined with model quality assessment is complementary to using larger sequence databases and a more extensive folding procedure.

**Table 4.** Number of Pfam families with unknown structure that can be modeled at 1% and 10% FPR of which the overlap with the Baker studies are given by number in parenthesis.

|  | 0.01 | 0.1 |
| --- | --- | --- |
| ProQ3D | 36 (10) | 225 (75) |
| PcombC | 42 (21) | 179 (74) |
| Pcons | 18 (9) | 218 (91) |
| CNS-contact | 62 (13) | 232 (38) |
| Union | 114 (35) | 558 (143) |
| All | 6379 (558) | 6379 (558) |

In Table 5 it can be seen that the average length of the successfully predicted models is shorter than for the average length of all models. The PconsC3 scores are also stronger, as expected. However the number of effective sequences and other properties are surprisingly not that different.

**Table 5.** Properties of Pfam families that can be modeled accurately at FPR 0.1. Statistical significant differences from a students t-test at P-values 0.01 and 10^−5^ are marked with * and ** respectively for all columns except the last.

|  | <PconsC3 score> | Helix | Sheet | Coil | <Meff> | <Length> | Transmembrane | <TM-score> |
| --- | --- | --- | --- | --- | --- | --- | --- | --- |
| Union | 0.41** | 0.33* | 0.18* | 0.5 | 427* | 104** | 0.11** | 0.56 |
| PfamPDB | 0.45** | 0.34 | 0.21** | 0.46** | 1075** | 126** | 0.04** | 0.53 |
| NoPDB | 0.32 | 0.36 | 0.15 | 0.50 | 300 | 187 | 0.25 |  |

Of the 558 models predicted at a FPR of 10% 143 have also been predicted by the Baker group. The comparison of these models is not absolutely trivial as different protein sequences have been used. However, in 74% of the cases our models have a TM-align score of 0.50 or higher indicating that they represent the same fold. In some cases where the models differs it appears as our models are mirror images of the Baker models.

Figure 6 shows a side-by-side comparison of PconsFold2 models (a) and (c) with those predicted by the Baker group (b) and (b) for two exemplary Pfam domains. PF02660 in Fig. 6 (a) and (b) is a Glycerol-3-phosphate acyltransferase transmembrane protein domain family. The TM-score between the two models is 0.79. It can be seen that PconsFold2 misses helical regions in the termini (red and blue ends of the model). The models for the Glycosyl transferase WecB/TagA/CpsF family of PF03808 in Fig. 6 (c) and (d) have an agreement of 0.75 TM-score. Again in the PconsFold2 model the secondary structural elements are not as defined as in the Rosetta model by the Baker group. It is to be noted though that the PconsFold2 models are taken directly from the output of CONFOLD stage 1 and have not been further refined.

**Fig. 6.**
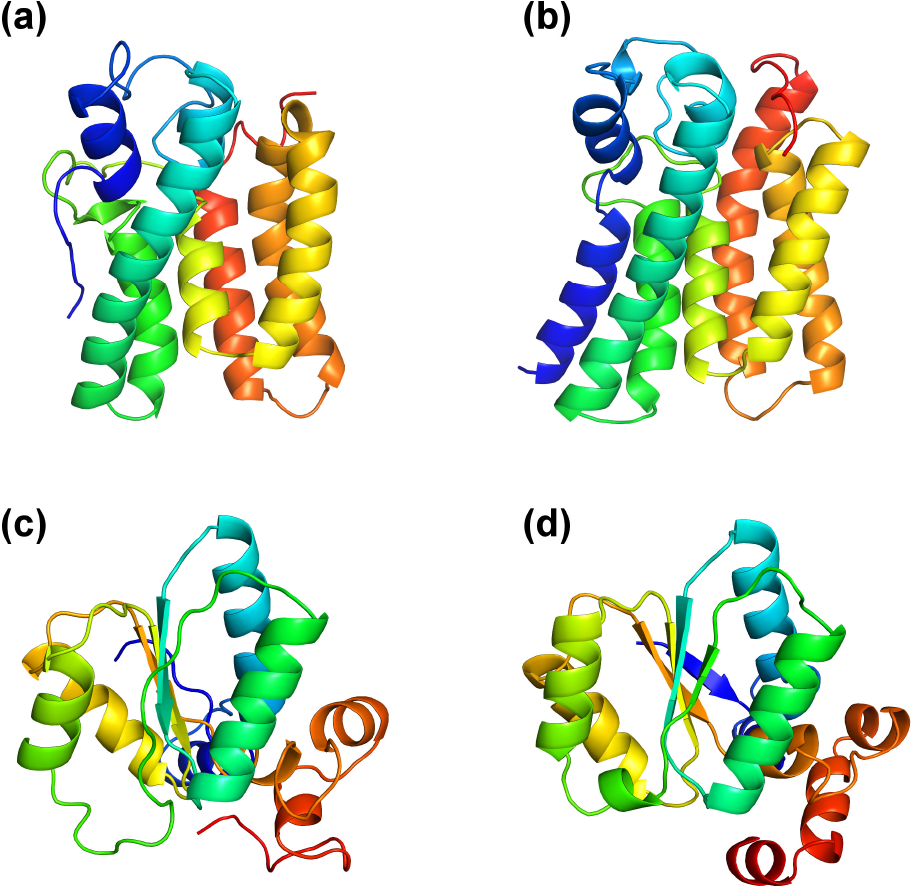
Comparison for two example Pfam families (a) PconsFold2 model for PF02660 (b) model by the Baker group for the same family (c) PconsFold2 model for PF03808 (d) model by the Baker group for the same family.

## 5 Conclusion

In this study we first present a novel protein folding pipeline, PconsFold2 that combines contact prediction, structure generation and model quality estimations. We show that the model quality estimation is an important step in the pipeline as the number of models that reliably can be predicted increases significantly when it is included. We also use this pipeline to predict the structure of 6379 Pfam families of unknown structure. At a FPR of 10% we find 558 Pfam families. The structure of 74% of these has not been reported before in any prediction study. Further, these models are obtained without the use of meta-genomic data, and the number of accurate models might therefore increase significantly if such sequences were included.

## Acknowledgements

We thank Björn Wallner, Christian Blau, and the entire CASP community for valuable discussions.

## Funding

This work has been supported by the grants from the Swedish Research Council (VR-NT 2012-5046) and Swedish e-Science Research Center. Computational resources were provided by the Swedish National Infrastructure for Computing (SNIC) at NSC, HPC2N and Uppmax.

